# RhoA/ROCK2 signalling is enhanced by PDGF-AA in fibro-adipogenic progenitor cells in DMD

**DOI:** 10.1101/2021.04.12.439417

**Authors:** Esther Fernández-Simón, Xavier Suárez-Calvet, Ana Carrasco-Rozas, Patricia Piñol-Jurado, Susana López-Fernández, Joan Josep Bech Serra, Carolina de la Torre, Noemí de Luna, Eduard Gallardo, Jordi-Díaz-Manera

**Affiliations:** Neuromuscular Diseases Unit, Neurology Department, Hospital de la Santa Creu i Sant Pau and Biomedical Research Institute Sant Pau (IIB Sant Pau), Barcelona, Spain; Centro de Investigaciones Biomédicas en Red en Enfermedades Raras (CIBERER), Madrid, Spain; Plastic surgery Department, Hospital de la Santa Creu i Sant Pau, Autonomous University of Barcelona, Barcelona, Spain; Proteomics Unit, Josep Carreras Leukaemia Research Institute, Barcelona, Spain; John Walton Muscular Dystrophy Research Center, University of Newcastle, UK

**Keywords:** Duchenne muscular dystrophy, Fibro-adipogenic precursor cells, Platelet-derived growth factor, Muscular dystrophies, Fibrosis

## Abstract

The lack of dystrophin expression in Duchenne muscular dystrophy (DMD) leads to muscle necrosis and replacement of muscle tissue by fibro-adipose tissue. Although the role of some growth factors in the process of fibrogenesis has been previously studied, the pathways that are activated by PDGF-AA in muscular dystrophies have not been described so far. Herein we report the effects of PDGF-AA on the fibrotic process in muscular dystrophies by performing a quantitative proteomic study in DMD isolated fibro-adipogenic precursor cells (FAPs) treated with PDGF-AA. In vitro studies showed that RhoA/ROCK2 pathway is activated by PDGF-AA and induces the activation of FAPs. The inhibition of RhoA/ROCK signalling pathway by C3-exoenzyme or fasudil attenuated the effects of PDGF-AA. The blocking effects of RhoA/ROCK pathway were analysed in the dba/2J-mdx murine model with fasudil. Grip strength test showed an improvement in the muscle function and histological studies demonstrated reduction of the fibrotic area. Our results suggest that blockade of RhoA/ROCK could attenuate the activation of FAPs and could be considered a potential therapeutic approach for muscular dystrophies.

## INTRODUCTION

Duchenne muscle dystrophy (DMD) is an X-linked recessive muscular disorder produced by mutations in the *DMD* gene (Birnkrant et al. 2018). Mutations in the *DMD* gene are associated to absent or very reduced dystrophin expression leading to a fragile muscle membrane susceptible to be damaged during muscle contraction (Wallace and McNally 2009). Muscle membrane fragility is the responsible of chronic injury resulting on continuous cycles of necrosis and regeneration of muscle fibres, persistent inflammation and eventually loss of muscle fibers and their replacement by fat and fibrous tissue with no contractile properties leading to permanent weakness and disability (Bushby et al. 2010; Klingler et al. 2012).

Fibro-adipogenic precursor cells (FAP) are mesenchymal progenitors muscle-resident cells characterized by the expression of platelet-derived growth factor receptor alpha (PDGFRα). FAPs are the main responsible of the fibrotic and fatty tissue expansion in pathologic conditions in the skeletal muscles (Joe et al. 2010; Perandini et al. 2018; Uezumi et al. 2011). It has been shown that different external stimulus can lead FAPs to differentiate into adipocytes or fibroblasts. FAPs are activated upon acute injury, proliferate and release different components of the extracellular matrix (ECM) to serve as scaffold where the new regenerated myofibers are displayed. After muscle regeneration, excessive FAPs are cleared by apoptosis mediated by tumour necrosis factor alpha (TNF-α)(Fiore et al. 2016). However, in dystrophic muscles, growth factors released by M2 macrophages continuously activate FAPs proliferation, that release collagen-I and other components of the ECM leading to an expansion of the fibrotic tissue(Perandini et al. 2018; Phelps, Stuelsatz, and Yablonka-Reuveni 2016). The molecular pathways driving FAPs fate in muscular dystrophies are just started to be understood. It is well-known that TGF-β promotes FAP proliferation, inhibits TNF-α mediated FAP apoptosis and drives FAPs to differentiate into fibroblasts. Concomitantly, TGF-β can also enhance the production and release of ECM components such as collagen or fibronectin. Once released, these matrix elements are the main component of the fibrotic tissue that is progressively accumulated in skeletal muscles of patients with muscular dystrophies (Kharraz et al. 2014; Mann et al. 2011).

Apart from TGF-β, other cytokines and growth factors have been shown to drive fibrosis accumulation such as connective tissue growth factor (CTGF) or platelet-derived growth factors (PDGFs). PDGF-AA has a prominent role in other fibrotic diseases, such as pulmonary fibrosis, skin or hepatic fibrosis (Olson and Soriano 2009). PDGF-AA is produced by different cells like platelets, endothelial cells or macrophages and acts via the PDGFRα (Scotton and Chambers 2007). PDGFRα is a tyrosin kinase receptor with a well characterized intracellular function. It is known that binding of ligands induces receptor dimerization and autophoshorylation of the intracellular domain, activating the kinase domain and providing docking sites for downstream signalling molecules (Kazlauskas and Cooper 1989; Kelly James D et al. 1991). Autophosphorylation of PDGFRα triggers different signalling pathways such as Ras-MAPK, PI3K or PLC-ƴ which are involved in different cellular responses. Examples of the cellular responses triggered by PDGF-AA signalling are proliferation, cellular differentiation, apoptosis inhibition, mobilization of intracellular calcium or cell motility (Andrae, Gallini, and Betsholtz 2008; Svegliati et al. 2007).

Although the role of PDGF-AA has been widely studied in fibrotic processes, its role in driving muscle fibrosis in muscular diseases is not completely known. Previous studies have shown that PDGF-AA expression is higher in dystrophic than in healthy muscles (Zhao et al. 2003). Several tyrosin-kinase inhibitors blocking the PDGFRα reduced muscle fibrosis in preclinical animal models of muscular dystrophy by inhibiting fibroblast proliferation, migration and reducing the release of ECM components(Makino et al. 2017, Piñol-Jurado et al., 2018; Ieronimakis et al. 2016).However, the pathways activated by PDGF-AA in FAPs in muscular dystrophies have not been described so far. This knowledge would allow us to identify new targets for the development of therapies aiming to reduce fibrosis in patients with these diseases and slow down the on-going degenerative process. In this study, we have used mass spectrometry to identify Rho-kinase as one of the pathways activated by PDGF-AA in FAPs isolated from human skeletal muscles. These results allowed us to demonstrate the utility of fasudil, a pan-rho-kinase inhibitor improving muscle strength and reducing muscle fibrosis in a murine model of DMD.

## MATERIAL & METHODS

### Cell culture

FAPs were obtained from 3 DMD patients muscle biopsy. To obtain FAPs from human muscle biopsies, fat and capillaries were removed in Hank’s Balanced Salt Solution medium (HBSS). Muscle explants were cultured and once the different cell types began to migrate from the explant, an immunomagnetic separation was done using specific anti-CD56 antibodies (MiltenyiBiotec, Bergisch-Gladbch, Germany). The negative fraction (CD56-) was cultured in Dulbecco’s Modified Eagle’s Medium (DMEM) and 199 (M199) medium (3: 1) supplemented with 10% fetal bovine serum (FBS),10 mM glutamine and penicillin-streptomycin (all from Lonza, Basel, Switzerland) and their purity was verified with the fibroblast marker TE-7 (Merck) by immunofluorescence (IF) and PDGFRα expression by fow cytometry. Representative images of the IF and flow-cytometry are shown in supplementary figure 1A and 1B. Stained cells were analysed by MACSQuant 10 (milteny Biotec). Compensations were adjusted according to the single stained controls. FlowJo v10 software was used for data analysis and data elaboration.

FAPs were cultured until obtaining a confluence of approximately 80% in proliferation medium with 10% FBS and then changed to a basal medium (DMEM: M199 (3:1) supplemented with 0.5% FBS, 10 mM glutamine and penicillin-streptomycin (all from Lonza)) for 24 hours. PDGF-AA was added to the medium at 50 ng/ml each day during a total of 4 days (Supplementary figure 1C). To test the activation of the RhoA pathway with PDGF-AA, the pathway was first inhibited with fasudil 50µM (Bio-Techne, Minnesota, USA) or C3-exoenzyme 2µg/ml (Cytoskeleton, Denver, USA) for 15 hours and then activated with PDGF-AA at 50ng/ml for 20 minutes and proceed to the G-LISA or IF assay of myosin light chain phosphorylation (p-MLC). To test the functional effect of the PDGF-AA pathway and its inhibition, C3-exoenzyme at 2µg/ml or fasudil at 50µM was first added to the culture and then PDGF-AA was added 4 hours later at 50 ng/ml. This inhibition/induction cycle was repeated for as many days as the test lasted. Four different conditions were tested: untreated FAPs, treated with C3-exoenzyme and PDGF-AA, with fasudil and PDGF-AA and with PDGF-AA alone.

### Protein extraction, quantification and digestion for quantitative proteomics

The protein samples were extracted with 6M Urea / 200mM Ammonium Bicarbonate (ABC) and precipitated with cold acetone. Cleaned samples were solubilized in the previous buffer described above and then quantified with the RCDC Protein Assay kit (Biorad, Hercules, CA). Ten μg of protein from each sample were digested in-solution using both Lys-C and Trypsin. Briefly, the samples were reduced with 10mM dithiothreitol (DTT, in 200mM ABC) for 1h at 30ºC and 650 rpm in the thermo-mixer, alkylated with 20mM chloroacetamide (CAA, in 200mM ABC) for 30 minutes at room temperature in the dark at 650 rpm in the thermo-mixer. Then, samples were diluted to 2M Urea final concentration and the required amount of 1μg/μl LysC (WAKO, Osaka, Japan) was added to obtain a 1:10 ratio enzyme:protein (w:w). The digestion was performed overnight at 37ºC at 650 rpm in the thermo-mixer. After that, samples were diluted at 1M Urea final concentration. Finally, the required amount of 1μg/μl trypsin (Promega, Madison, WI) was added to obtain a 1:10 ratio enzyme:protein (w:w) and incubated for 8h at 37ºC at 650 rpm in the thermo-mixer. Peptide mixtures were desalted using the commercial columns Ultra Microspin C18, 300A silica (The Nest Group, MA, USA) according to the manufacturer instructions. Finally, the samples were dried in a SpeedVac and kept at -20ºC until the LC-MS/MS analysis.

### LC-MS/MS analysis

The peptide samples were analysed using a Lumos Orbitrap mass spectrometer (Thermo Fisher Scientific, San Jose, CA, USA) coupled to an EASY-nLC 1000 (Thermo Fisher Scientific (Proxeon), Odense, Denmark). Peptide were loaded directly onto the analytical column and were separated by reversed-phase chromatography using a 50cm column with an inner diameter of 75 μm, packed with 2 μm C18 particles spectrometer (Thermo Fisher Scientific) with a 110 min run, comprising consecutive steps with linear gradients from 5% to 22% B in 80 min, from 22% to 32% B in 10 min and from 32% to 95% B in 20min at 250 nL/min flow rate and a binary solvent system of 0.1% formic acid in H2O (Solvent A) and 0.1% formic acid in acetonitrile (Solvent B). After each analysis, the column was washed for 10 min with 5% buffer A and 95% buffer B.

The mass spectrometer was operated in a data-dependent acquisition (DDA) mode and full MS scans with 1 micro scans at resolution of 120.000 were used over a mass range of m/z 350-1500 with detection in the Orbitrap. Auto gain control (AGC) was set to 2E5 and dynamic exclusion to 60 seconds. In each cycle of DDA analysis, following each survey scan Top Speed ions with charged 2 to 7 above a threshold ion count of 1e4 were selected for fragmentation at normalized collision energy of 28%. Fragment ion spectra produced via high-energy collision dissociation (HCD) were acquired in the Ion Trap, AGC was set to 3e4, isolation window of 1.6 m/z and maximum injection time of 40 ms was used. All data were acquired with Xcalibur software v3.0.63.

### Raw data processing and database search

The samples were analysed with the MaxQuant software (version 1.6.1.0) through the human Swissprot database. Trypsin was chosen as enzyme and a maximum of two missed cleavages were allowed. Carbamidomethylation (C) was set as a fixed modification, whereas oxidation (M) and acetylation (N-terminal) were used as variable modifications. Searches were performed using a peptide tolerance of 10 ppm and a product ion tolerance of 0.5 Da. Resulting data files were filtered for FDR <1% at both peptide and protein level. The non-unique peptides were assigned to the corresponding protein group according to the Razor peptides rule implemented in the software. The option “match between runs” was also enabled. The intensity values were normalized using the LFQ algorithm.

### Statistical and Bioinformatic analysis

The final list of proteins was analysed using R. First, the protein list was filtered to remove the peptides/proteins tagged as “Reverse” (significantly identified in the reverse database), “potential contaminant” (items identified as contaminants in the “contaminants.fasta” file) and “Only identified by site” (proteins identified only with modified peptides). Then, we selected the proteins with at least 75% of valid values in each experimental condition. The missing values were imputed using the knn algorithm. The comparisons between groups were done using a t-test. The p-values were adjusted using the Benjamini-Hochberg procedure.

The total proteins analysed were represented in a volcano plot generated by Graphpad Prism Software. Proteomap was performed with upregulated proteins in PDGF-AA treated cells and was generated to visualise the differential contribution of biological pathways (bionic-vis.biologie.uni-greifswald.de/, v2.0). The map is created from the Kyoto Encyclopedia of Genes and Genomes (KEGG) functional classification and it shows the quantitative composition of proteomes. Each protein is represented by a polygon: areas reflect protein abundance, and functionally related proteins are arranged in common and similarly colored regions (Liebermeister et al. 2014). The area for each protein reflects the magnitude of the fold change in PDGF-AA treated cells in comparison with untreated cells. The upregulated proteins represented in the proteomap were also analysed for enrichment of pathways in the Reactome database and were represented in a heatmap generated by the Clustvis online tool (biit.cs.ut.ee/clustvis/)(Fabregat et al. 2018).

### Cell viability

The effect of each treatment in FAPs viability was measured with the PrestoBlue reagent (Invitrogen). Cells were seeded into 96-well plates at 80% confluence and incubated in the presence or absence of each of the drug treatments for either 30 min, 24, 48 or 72 hours period. Cell viability was measured according to the manufacturer’s instructions. Fluorescence was measured at wavelengths 570 nm excitation and 600 nm emission using a microplate reader (INFINITE M1000 PRO, Tecan Trading AG, Switzerland).

### G-LISA

To test whether PDGF-AA promoted the RhoA pathway in DMD FAPs, the RhoA-GTP protein was analysed by a colorimetric assay. Cells were seeded into 96-well plates at 80% confluence and the medium was changed to 0.5% FBS basal medium for 24 hours. The pathway was inhibited with the C3-exoenzyme (Cytoskeleton) inhibitor at 2 µg/ml overnight. Then, the pathway was activated with PDGF-AA at 50 ng/ml for 20 minutes at 37°C and the G-LISA assay (Cytoskeleton) was performed following the manufacturer’s instructions. The RhoA-GTP measurement was normalized to the amount of total RhoA that was measured by Western-blot.

### Proliferation Assay

The proliferation assay was performed with FAPs obtained from 3 different DMD patients and 3 replicates were used for each condition. FAPs were cultured in 96-well plates and 5,000 cells were seeded per well. Once cells were plated, media was changed to basal medium at 0.5 % FBS and 24 hours before, cells were treated with 50µM fasudil) or 2µg/ml C3-exoenzyme and PDGF-AA was added 4 hours later at 50 ng/ml. This inhibition/induction cycle was repeated as many days as the test lasted. A total of 4 different conditions were performed: untreated FAPs, treated with c3-exoenzyme and PDGF-AA, with fasudil and PDGF-AA and with PDGF-AA alone. The proliferation assay was carried out following the manufacturer’s instructions (Roche, Indianapolis, IN) at 24 and 72 hours. The result was measured using the Coulter AD 340 plate reader (Beckam-Coulter, Brea, CA, USA) with the AD-LD program.

### Migration assay

The migration assay was performed with FAPs obtained from 3 different DMD patients and 3 replicates were made for each condition. Cells were plated in 24-well plates with inserts separating 2 chambers (Ibidi, Munich, Germany). A total of 15,000 cells were seeded in each part of the insert and after 24 hours of culture in basal medium at 0.5% FBS, FAPs were treated with 25 µg/ml of mitomycin C (Sigma-Aldrich, San Luis, MO, USA) for 1 hour. Then, the insert was removed leaving a space of 500 µm between the 2 chambers and cells were treated with 50µM fasudil or 2µg/ml C3-exoenzyme and 4 hours later PDGF-AA was added at 50 ng/ml. This inhibition/induction cycle was repeated for as many days as the test lasted. At 48 and 72 hours, cells were fixed in 4% paraformaldehyde (PFA) and stained with Hoechst 33342. A total of 3 images were made at 10X of each well and the quantification of cells that migrated to the 500 µm space was quantified with Fiji software.

### Sircol

Total soluble collagen released by cultured FAPs was measured using a colorimetric assay. FAPs were seeded at 10,000 cells/cm^2^ in 24-well plates and after 24 hours, the medium was changed to basal medium at 0.5% FBS. A total of 4 inhibition/induction cycles were performed and the test was carried out with the commercial Sircol kit, following the manufacturer’s instructions (Biocolor, UK). The result was measured using the Coulter AD 340 plate reader (Beckam-Coulter, Brea, CA, USA) with the AD-LD program. At the end of the assay, the nuclei were stained with TO-PRO-3 (Thermo Fischer Scientific) and the signal emitted was measured with the Odyssey reader (LI-COR)(Supplementary figure 1E). The results obtained with the Sircol assay were normalized using the signal emitted by TO-PRO-3 to subtract the effect of the increased proliferation after PDGF-AA treatment.

### F-actin

The content of filamentous actin (F-actin) was assessed in cultured FAPs and 4 experimental conditions were performed: untreated, inhibited with C3-exoenzyme or 50µM fasudil and then activated with PDGF-AA for 20 minutes at 37 °C or only activated with PDGF-AA. After that time cells were fixed with 4% PFA, incubated with Rhodamine Phalloidin Reagent (Abcam, Cambridge, UK) for 1 hour and then stained with Hoescht 33342.

### Western-Blot

Cells corresponding to each condition or muscle tissue were lysed in RIPA buffer (Sigma-Aldrich) with protease and phosphatase inhibitors (Roche, Basel, Switzerland). Cell lysates were centrifuged at 4 °C at 13,000xg for 20 minutes and supernatants were stored at -80 °C. Protein concentration was quantified using the Pierce ™ BCA Protein Assay kit (Thermo Fisher Scientific). 30 µg of total protein was loaded onto a 10% polyacrylamide gel and transferred to a nitrocellulose membrane. Nonspecific binding sites were blocked for 1 hour in casein diluted 1: 1 with Tris-buffered saline (TBS). The membranes were incubated overnight with the primary antibody corresponding to each experimental condition. Corresponding secondary antibodies bound to Dye680 or Dye800 (Li-Cor) were incubated for one hour at 1: 7,500 dilution (table 1). The specific bands corresponding to the proteins of interest were visualized with the Odyssey infrared detection system (Li-Cor) and with the Image Studio program (Li-Cor). Total protein bands were measured with the Revert 700 kit (Li-Cor) (Supplementary figure 1D). The primary and secondary antibodies used in the study are listed in table 1.

**Table 1:** Primary, secondary antibodies and dyes used in this study. ^1^It is not an antibody.^2^Volume per test following manufacturer’s instructions. N/A: not applicable; Ms: Mouse; IHQ: Immunohistochemistry; FC: Flow cytometry. MACS: Magnetic activated cell sorting

| Target | Host | Dilution | Application | Manufacturer |
| --- | --- | --- | --- | --- |
| ArHGEF2 | Rabbit | 1/100 | WB | Sigma-Aldrich |
| RhoA | Rabbit | 1/100 | WB | Cell Signalling |
| p-MLC2 | Mouse | 1/100 | IF | Cell Signalling |
| PDGFRα | Goat | 1/50 | IF | R&D System |
| F4/80 | Rabbit | 1/200 | IF | R&D System |
| Wheatgerm agglutinin-488 | N/A | 1/100 | IF | Invitrogen |
| Hoescht 33342 | N/A | 1/1000 | IF | Invitrogen |
| CD56 | Ms | N/A | MACS | Miltenyi Biotech |
| PDGFRα-biot | Goat | 1/75 | FC | R&D System |
| CD56-PE | Ms | 1/20 | FC | Miltenyi Biotech |
| LIVE/DEAD<sup>1</sup> | N/A | 1/1000 | FC | Thermo Fisher |
| F4/80 | Rat | N/A | FC | Bio Legend |
| CD45 | Rat | N/A | FC | eBioscience |
| Integrin- α7 | Rat | N/A | FC | Milteny Biotec |
| Ter119 | Rat | N/A | FC | eBioscience |
| CD31 | Rat | N/A | FC | Thermo Fisher |
| Sca1 | Rat | N/A | FC | BD |
| CD163 | Rat | N/A | FC | eBioscience |
| CD206 | Rat | N/A | FC | Bio Legend |
| CD80 | REA | N/A | FC | Milteny Biotec |
| HLA-DR | Rat | N/A | FC |  |

| Secondary | Dilution | Conjugation | Application | Manufacturer |
| --- | --- | --- | --- | --- |
| Donkey anti-goat | 1/500 | Alexa594 | IF | Invitrogen |
| Donkey anti-rabbit | 1/500 | Alexa647 | IF | Invitrogen |
| Goat anti-mouse | 1/500 | Alexa488 | IF | Invitrogen |
| Goat anti-rabbit | 1/7500 | IRDye 800 | WB | LICOR |
| Goat anti-rabbit<br>biotinilyated | 1/2000 | N/A | WB | Invitrogen |
| Streptavidin 800 | 1/2000 | IRDye 800 | WB | LICOR |
| Streptavidin PE-Cy5 | 1/200 | PE-Cy5 | FC | Biolegend |

### Histology and Immunofluorescence

Cultured FAPs were fixed with ethanol and incubated with blocking solution (Santa Cruz Biotech, Dallas, TX, USA)). Frozen muscle sections were obtained with a cryostat (Leica Microsystems, Wetzlar, Germany), fixed with acetone and incubated with blocking solution (Santa Cruz Biotech). The primary and secondary antibodies used in the study are listed in table 1.

Images were obtained with an Olympus BX51 microscope coupled to an Olympus DP72 camera. Image J software was used to quantify the intensity of individual cells and to quantify the area of positive staining in muscle tissue. A minimum of five independent fields per staining and individual cells were quantified(Schindelin et al. 2012).

### Mouse model

Animal procedures were performed according to the National Institutes of Health Guidelines for the Care and Use of Laboratory Animals and were approved by the Ethical Committee of the Universitat Autònoma de Barcelona. The animals used in this study were all 7-weeks old and males. A total of 6 DBA/2J-mdx males were treated with fasudil (LC Laboratories) at 100 mg/kg/day orally for 6 weeks. Five untreated DBA/2J-mdx males were used as controls and 5 DBA/2J males were used as healthy mice.

### Grip strength test

Force was measured using a Grip Strength Meter. Mice animals were initially held by the tail above the grid. Once mice grasped the grid, they were pulled out until release. The procedure was repeated 5 times and the mean of the 3 highest values was used. The mean result was normalized to the mice weight. Measurements were carried out using a sensor. Readings were expressed in newtons (N) and normalized to the body weight of the animal. Results are presented as grip force/weight (N/g).

### Flow cytometry

Single cell suspension from cell culture, quadriceps and tibial anterior were obtained for flow cytometry. Cultured human muscle derived cells were labelled using anti-PDGFRα-followed by streptavidine-PECy5 and anti-CD56. Cells obtained from mice were labelled using anti-sca1, anti-CD45, anti-F4/80, anti-CD163 and a viability marker (table 1). Samples were acquired with the MACSQuant Analyzer 10 flow cytometer (MiltenyiBiotec). Doublet cells were excluded using Forward scatter area and height. Compensations were adjusted according to the single stained controls. Total viable FAPS (Viable, Ter119-/CD45/CD31-/integrin-α7-/sca1+) and macrophages (Viable, CD45+ F4/80+) counts were quantified using MACSQuantify software and normalized according to grams of muscle tissue. FlowJo v10 software was used for data analysis and data elaboration.

### Statistical Analysis

Results are expressed as mean ± standard error of means (SEM). Differences among the groups were analysed using one-way ANOVA Test. When ANOVA revealed significant differences, the Tukey post hoc test was performed. The significance level was set at p<0.05. The statistical analyses were calculated using GraphPad Prism version 8 for Windows (San Diego, CA: GraphPad Software, Inc)

## RESULTS

### PDGF-AA induces activation of Rho-A pathway in human muscle FAPs

We treated FAPs isolated from muscle biopsies from 3 DMD patients with 50 ng/ml of PDGF-AA for 4 days. In order to analyse changes in the protein profile, we used quantitative proteomic analysis with mass spectrometry comparing PDGF-AA treated FAPs against non-treated FAPs. We observed that 1890 proteins were differentially expressed (Volcano plot in figure 1A). We then constructed a proteomap of the upregulated proteins according to the functional gene classification (Kyoto Encyclopedia Genes and Genomes (KEGG)) (Figure 1B). In summary we observed an increase in the expression of proteins involved in various signalling pathways (MAPK, PI3K-Akt, FoxO or Hedgehog), cell metabolism and genetic information processing (Figure 1C-D). Among the different protein groups that were upregulated, we were interested in those proteins involved in the cell functions related to the fibrogenic process such as cytoskeletal rearrangement, cell adhesion, and rearrangement of actin filaments. We observed that proteins related to these cellular functions, such as proteins of the Ras homolog gene pathway (Rho), were increased in FAPs treated with PDGF-AA as displayed in a heatmap showing the differences between DMD FAPs untreated and treated with PDGF-AA (Figure 1E). Among the different proteins of this family, we focused on the Ras homolog gene family member A (RhoA) pathway since it has been involved in fibroblast activation and fibrosis in several other pathologies (Alkasalias et al. 2017)(Ge et al. 2016)(Datta et al. 2017a).

**Figure 1.**
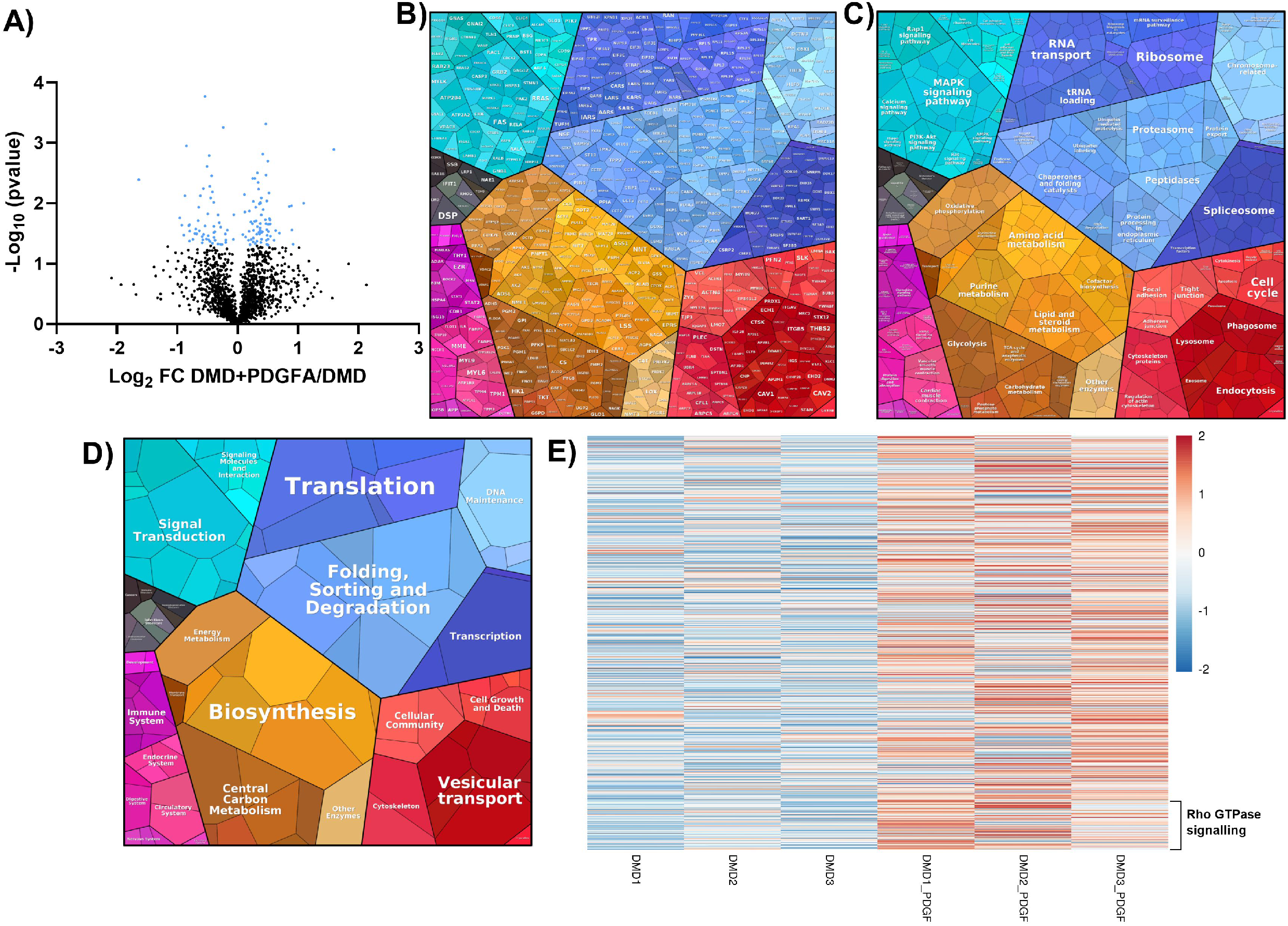
Proteome analysis of PDGF-AA treated DMD FAPs. A) Volcano plot representing the total proteome. Data is presented as the protein abundance changes of PDGF-AA treated FAPs relative to untreated FAPs. Significantly differentially expressed proteins (student T test, p≤ 0.05) are highlighted in blue. B to D) Proteomemap of upregulated proteins after PDGF-AA treatment. The proteomic map visualizes the composition of the proteomes in terms of abundance and function of the proteins. Each protein is represented by a polygon, and the area of each polygon reflects the abundance of the proteins (calculated with fold-change). Functionally related proteins appear in adjacent regions. The three panels represent three hierarchy levels, being the first one the individual protein detected (B), the second one shows proteins grouped into pathways (C) and the last one shows the functional category associated to a group of proteins (D). E) Upregulated proteins shown in proteomaps were also represented in heatmap showing the results of Reactome database.

### C3 exoenzyme and Fasudil treatment block Rho-Kinase pathway activation mediated by PDGF-AA

We validated mass spectrometry results in two different experiments. First, we studied the expression of guanine nucleotide exchange factor Rho 2 (ArhGEF2), which is the effector protein in the Rho-kinase pathway, in muscle samples from DMD patients and aged and sex matched controls. We observed a 7.7-fold increase in the expression of ArhGEF2 in DMD muscles compared to control muscles (Figure 2A and B). In a second step we studied the effect of PDGF-AA in different components of the Rho-kinase pathway (figure 2A). We analysed if the addition of PDGF-AA at 50 ng/ml to DMD FAPs in culture induced a significant increase in RhoA bound to guanosine triphosphate (GTP) (RhoA-GTP) compared to untreated FAPs (c-). To further confirm whether the increase in RhoA-GTP was mediated via activation of RhoA pathway, the effects of pre-treatment with the exoenzyme C3 (C3) was also analysed. Exoenzyme C3 is an ADP ribosyl transferase that selectively ribosylates RhoA proteins at asparagine residue 41, rendering it inactive. As expected, PDGF-AA treatment induced a significant 1.88-fold increase in the RhoA-GTP levels while pre-treatment with C3 at 2 µg/ml blocked the effect induced by PDGF-AA (Figure 2C).

**Figure 2.**
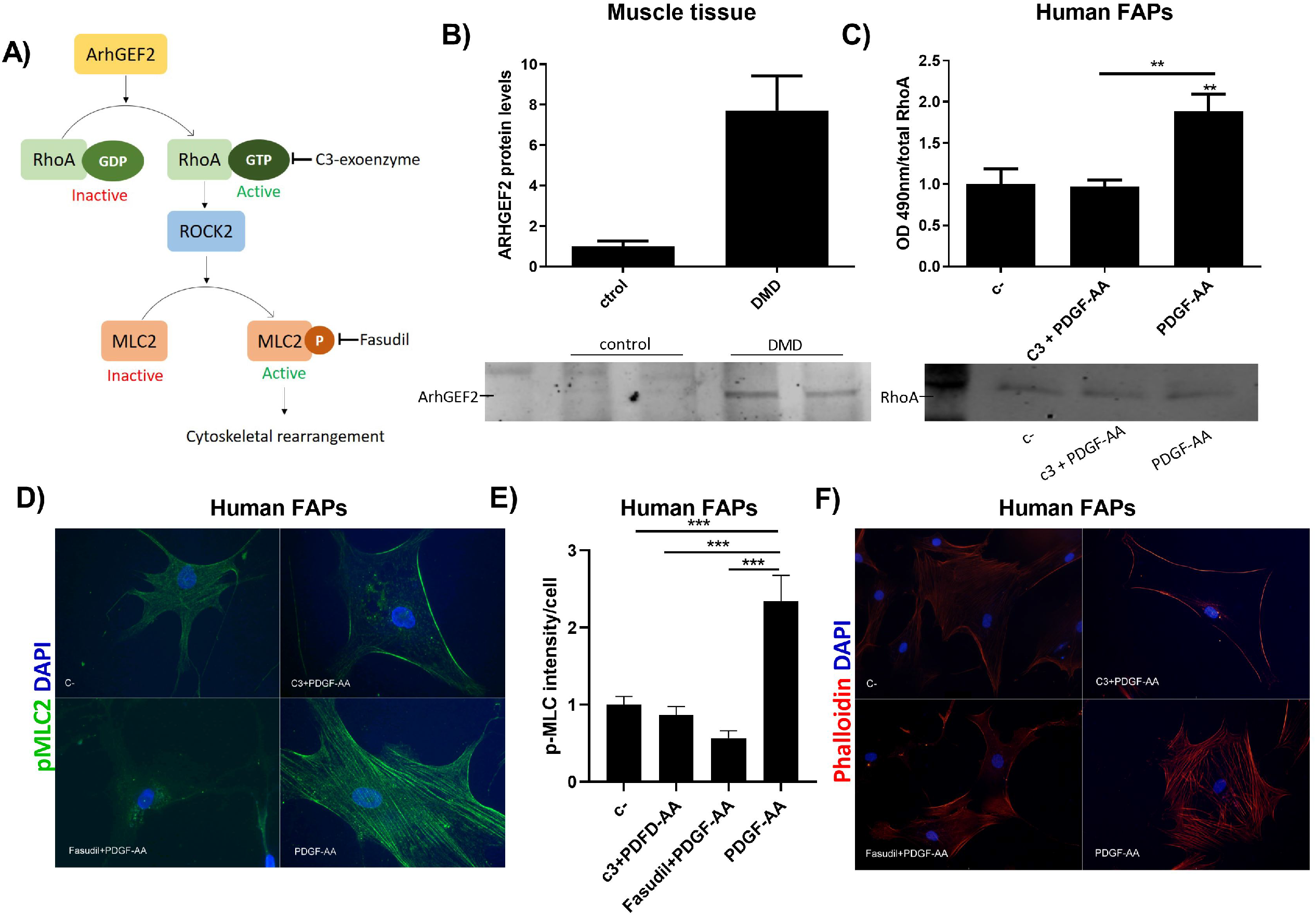
RhoA pathway is upregulated in DMD and PDGF-AA activates it. A) Molecular pathway of RhoA/ROCK2 signalling downstream of ArhGEF2 which catalyses the replacement of Rhoa-GDP to RhoA-GTP, controlling the RhoA activation. RhoA bound to GTP activates the Rho-associated coiled coil-forming protein kinase (ROCK), a serine/threonine kinase that regulates actin filament remodelling trough light chain myosin (MLC) phosphorylation. B)Quantification of WB bands of ArhGEF2 relative to total protein detection and representative bands of ArhGEF2 expression in muscle tissue obtained from healthy controls (n=2) and DMD patients (n=2). Total protein detection was used as loading control signal (Supplementary figure 1D). C) Optical density at 490 nm indicating RhoA activity relative to total RhoA protein detected by WB.D) Representative images of p-MLC staining using fluorescence microscopy. Immunofluorescence images were taken at 40X magnification. E) Quantification of p-MLC intensity of each cell after different treatments. F) Representative images FAPs stained with phalloidin after different treatments. Immunofluorescence images were taken at 40X magnification. C-: untreated FAPs. Data are represented as the mean of 3 replicates ± standard error of the mean. Results were statistically analysed using one-way ANOVA followed by Tukey post hoc test. Statistical significance was set at p<0.05. **p<0.01; ***p<0.0001.

Fasudil is a commercially available vasodilator used in the treatment of cerebral vasospasm also useful as treatment of pulmonar hypertension. Fasudil inhibits RhoA pathway by blocking the Rho-associated protein kinase (ROCK) which phosphorylates myosin light chain 2 (p-MLC2). Treatment of DMD FAPs with 50 ng/ml PDGF-AA promoted a statistically significant 2.3-fold increase in MLC2 phosphorylation, which was reversed by the pre-treatment of both C3-exoenzyme or fasudil as shown in Figure 2D and 2E. The last step of the Rho-kinase pathway is the polymerization of F-actin filament mediated by MLC2 phosphorylation. We observed that treatment of FAPs with PDGF-AA clearly induced polymerization on intracellular actin filaments, which was impinged by pre-treatment with C3 or fasudil (Figure 2F).

### Fasudil blocks PDGF-AA mediated increase in proliferation, migration and expression of collagen-I by human FAPs in vitro

We studied the effect of PDGF-AA, C3-exoenzyme and fasudil in proliferation, migration, and collagen-I production *in vitro*. Addition of PDGF-AA to the culture medium significantly increased FAPs proliferation at both 48 and 72 hours (3.6-fold and 6-fold increase, respectively) compared with untreated FAPs. In contrast, treatment of FAPs with C3-exoenzyme and fasudil prior to stimulation with PDGF-AA showed a significant decrease in proliferation rate (3.27-fold and 2.6-fold decrease, respectively) compared to FAPs treated with PDGF-AA at 72 hours (Figure 3A). All treatment doses and the different times of cell exposure were tested for cell viability. PrestoBlue data showed that C3-exoenzyme and fasudil had no effect on cell viability (Supplementary figure 1F).

**Figure 3.**
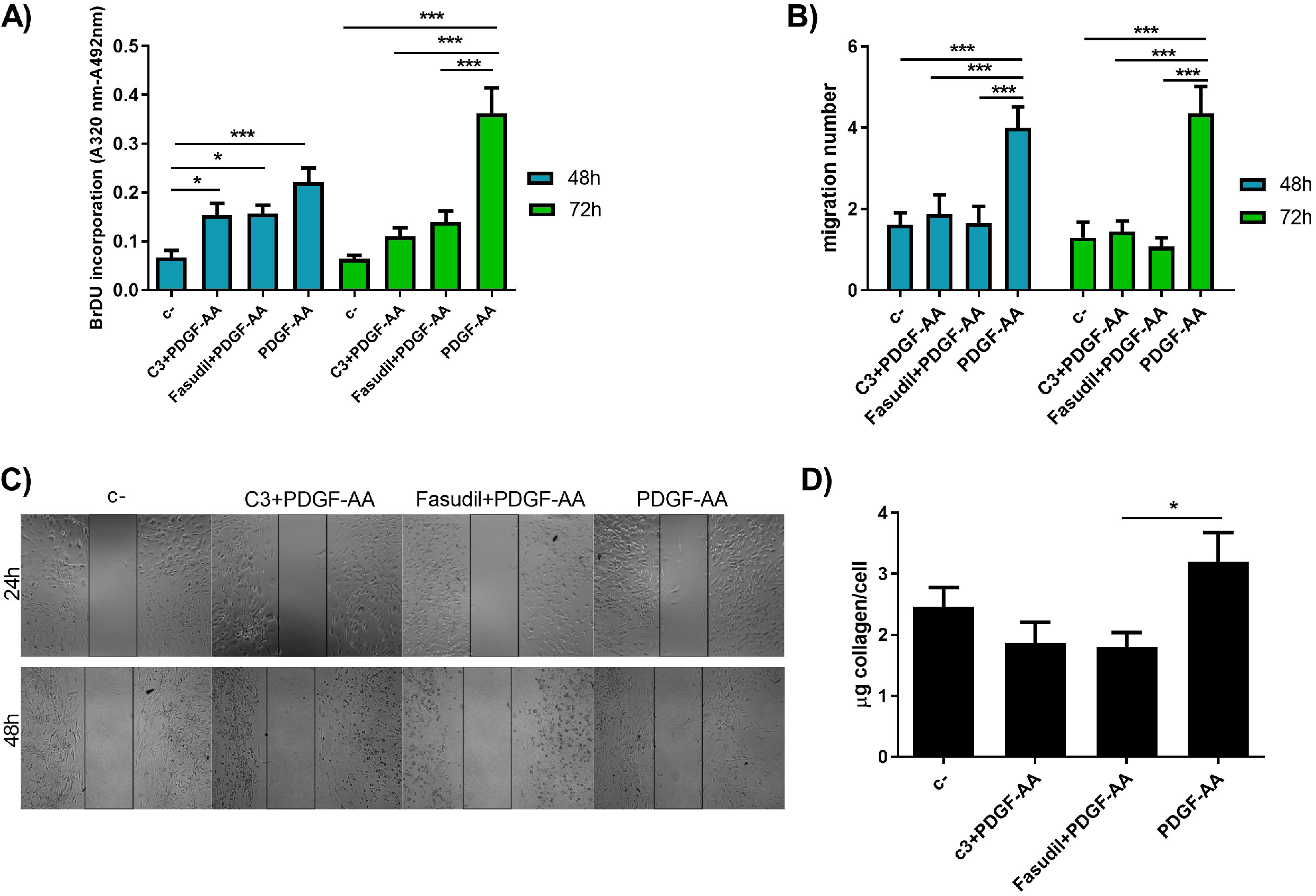
Effects of C3, fasudil and PDGF-AA on FAPs from DMD patients. A) FAP proliferation was analysed after 48 and 72 hours. B) FAP migration was analysed after 48 and 72 hours. C) Representative images of migration assay at both 48 and 72 hours. Images were taken at 5X magnification. D) The amount of collagen delivered to the supernatant was analysed after 4 days treatment. C-: untreated FAPs. Data are represented as the mean of 3 replicates ± standard error of the mean. Results were statistically analysed using one-way ANOVA followed by Tukey post hoc test. Statistical significance was set at p<0.05. *p<0.05; **p<0.01; ***p<0.0001.

Additionally, treatment with 50 ng/ml PDGF-AA significantly enhanced FAPs migration *in vitro* compared to untreated FAPs at both 48 and 72 hours (2.48-fold and a 3.37-fold increase respectively). This effect was significantly blocked when C3-exoenzyme and fasudil were added to the medium (Figure 3B-C) either at 48 or 72 hours.C3-exoenzyme decreased the migration rate by a 2.1-fold decrease at 48 hours and a 3-fold decrease at 72 hours. Fasudil decreased the migration rate by a 2.43-fold decrease at 48 hours and a 4-fold decrease at 72 hours.

PDGF-AA treatment also increased non-significantly the release of collagen-I to the culture medium an effect that was reversed by the addition of C3-exoenzyme and fasudil (Figure 3D). C3-exoenzyme significantly reduced the delivery of collagen-I by a 1.7 fold decrease.

### Fasudil treatment improves muscle function and reduces muscle fibrosis in the dba/2J-*mdx* murine model of DMD

Based in our *in vitro* results we decided to test if fasudil had an antifibrotic effect *in viv*o in a murine model of DMD. We treated 6 dba/2J-*mdx* mice of 7 weeks old with fasudil at a dose of 100 mg/kg/day orally for 6 weeks. At the end of the treatment, we analysed the effect of fasudil on animal weight and did not observe significant differences when compared with untreated mice (Supplementary figure, 2A). We also analysed muscle strength using forelimb grip-strength and compared the results with the ones obtained with non-treated 13 weeks old dba/2J-*mdx* mice and with 13 weeks healthy control mice. We observed a significant 1.76-fold increase in muscle strength of the forelimb in the treated mice compared with the non-treated mice (Figure 4A). Interestingly, differences between treated mice and WT mice did not reach significance. We also analysed the effect of fasudil in the quadriceps histology. Fasudil induced a significant 23% reduction in the collagen-I expression area in the quadriceps in the treated mice(Figure 4B and C). Additionally we observed a 42% decrease in the PDGFRα area a measure of the number of FAPs present in the tissue, in quadriceps of treated mice compared to non-treated *mice*, although these differences did not reach statistical significance (Figure 4 D and F). We also studied if fasudil had any effect in the heart of dba/2J-*mdx*. We did not see any change in the amount of fibrotic tissue in heart of the dba/2J-*mdx* at 13 weeks compared to the dba/2J-WT and consequently we did not observe any differences induced by the fasudil (Supplementary figure 2B and C).

**Figure 4.**
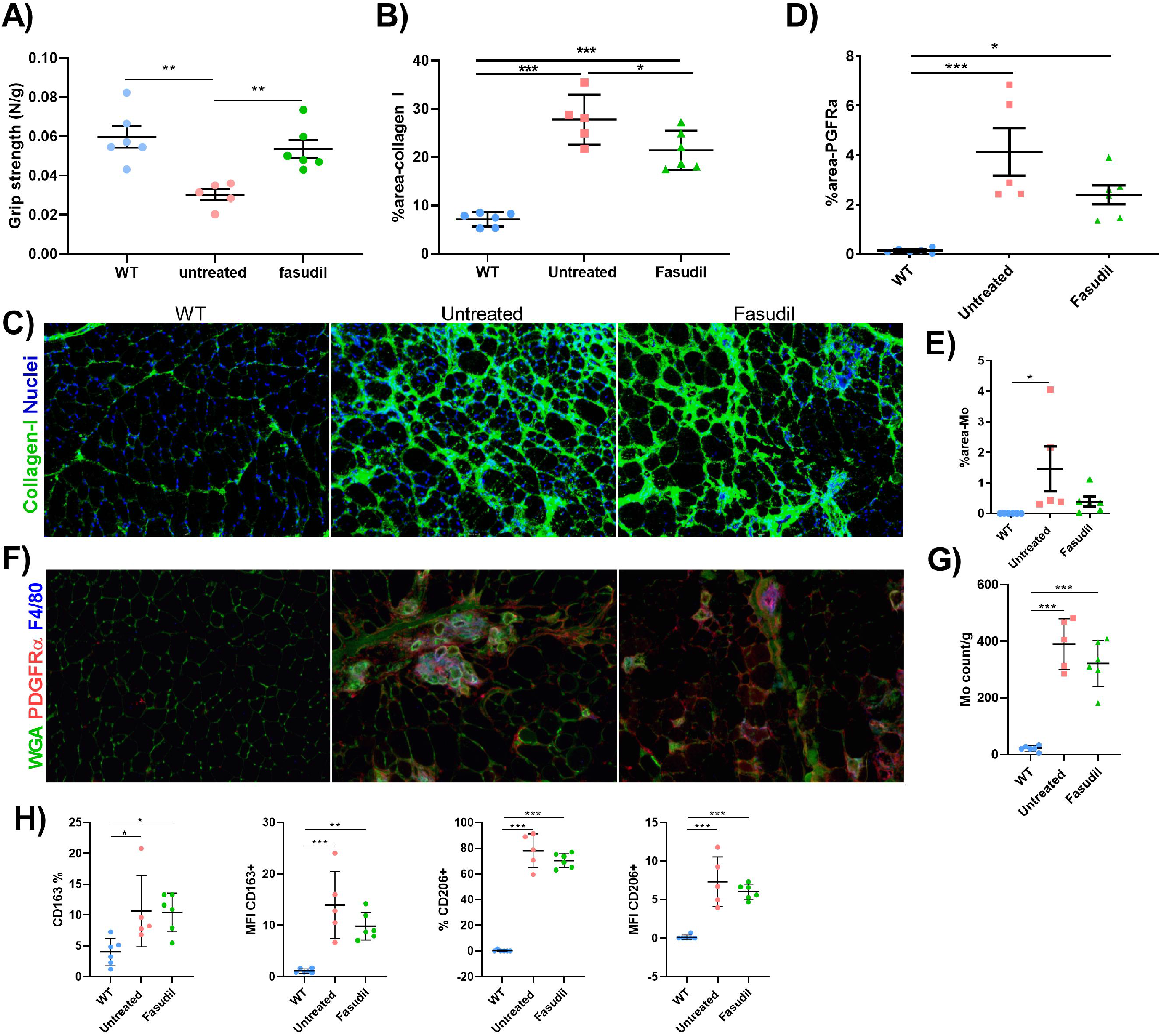
Effect of fasudil treatment in the dba/2J-mdx mice. A) Gripstrength test after 6 weeks treatment with fasudil in dba-2J WT mice, untreated dba/2J-mdx and fasudil treated dba/2J-mdx. Results are relative to weight of animals. B) Quantification of collagen I expression area in quadriceps. Immunofluorescence images were taken at 10X magnification. C) Representative immunofluorescence images of collagen I expression in muscle samples. D) Quantification of PDGFRα+ expression area in quadriceps. E) Quantification of macrophages expression area in quadriceps. F) Representative immunofluorescence images of PDGFRα+ (red), F4/80 (blue) expression in muscle samples. ECM is stained with wheat germ agglutinin (WGA) in green. Immunofluorescence images were taken at 10X magnification. G) Quantification of results obtained with flow cytometry to analyse the F4/80 macrophages (Mo) accumulated in quadriceps and tibial anterior obtained from mice. Data is represented as counts of F4/80 relative to the weight of quadriceps and tibial anterior. H) Percentage of total macrophages and mean fluorescence intensity (MFI) of CD163 and CD206 markers were analysed by flow cytometry. Data are represented as the mean of the groups analysed ± standard error of the mean. Results were statistically analysed using one-way ANOVA followed by Tukey post hoc test. Statistical significance was set at p<0.05. *p<0.05; **p<0.01; ***p<0.0001.

### Effect of fasudil treatment in inflammation

Quantification of macrophages was performed by analysing the area of the tissue positive for the F4/80 maker in quadriceps. Fasudil induced a 74% reduction in treated mice when compared to untreated mice (Figure 4F and E). Those results were in accordance with flow cytometry analysis which showed a 18.3% decrease in the number of F4/80 positive cells in treated mice compared to non-treated mice (Figure 4G). Indeed, we observed a non-significant trend towards a reduction in the expression of profibrotic M2 markers, such as CD163 and CD206, in macrophages obtained from fasudil treated mice. In detail, we observed a 31% reduction in the expression levels of CD163 marker in macrophages obtained from skeletal muscles of treated mice although there was only a 3% less number of CD163+ cells compared to non-treated mice. Similarly, we observed a 18% reduction in the expression levels of CD206 marker in macrophages obtained from skeletal muscles of treated mice although there was only a 10% less number of CD206+ cells compared to non-treated mice (Figure 4H). On the other hand, we did not observe any difference either in the expression levels or the number of CD209 positive cells, another M2 marker. We did not observe changes in the percentage of positive cells or in the expression of CD80 and DR, two well-known M1 markers, in the macrophages obtained from skeletal muscles of treated mice compared to non-treated mice (Supplementary figure 2D).

## DISCUSSION

In the present study, we have investigated the molecular pathways activated by PDGF-AA in skeletal muscles of patients with DMD driving to muscle fibrosis. We have observed that PDGF-AA activates RhoA/ROCK2 pathway which can be effectively blocked by fasudil, a well-known Rho-kinase inhibitor that reduces FAPs activation *in vitro*. We performed a proof-of-concept preclinical study to validate these findings and we observed a reduction in muscle fibrosis in the murine model of DMD. Our results expand the molecular pathways that are involved in the process of muscle fibrosis in DMD and support further investigations in the efficacy of Rho-kinase inhibitors as a treatment of DMD in particular and muscular dystrophies in general.

Absence of dystrophin in DMD produces sarcolemma instability leading to injury of muscle fibres that undergo continuous cycles of necrosis and repair leading to expansion of fibrotic and adipogenic tissue (Hogarth et al. 2019), (Perandini et al. 2018). Different growth factors have been related to the fibrotic expansion, however the role of PDGF-AA has not been fully studied. In order to understand the molecular consequences of persistent FAPs exposure to PDGF-AA, we treated FAPs from DMD patients with PDGF-AA for 4 days and observed an upregulation of several proteins belonging to the RhoA/ROCK2 pathway. The RhoA/ROCK2 signalling pathway is well known for regulating actin cytoskeleton organization and cellular dynamics in several cell types. RhoA acts through two molecular conformations: it is inactive when bound to a guanosine diphosphate (GDP), and active when bound to GTP. ROCK2, the Rho downstream effector molecule, is a serine/threonine kinase protein that regulates actin filament remodelling by phosphorylating numerous downstream target proteins, including the myosin binding subunit of myosin light chain 2 (MLC2) phosphatase (Chen, Pavlish, and Benoit 2008; Shang et al. 2013.). The actin reorganization triggered by RhoA/ROCK2 signalling pathway has demonstrated to drive fibroblasts activation in different pathologic conditions such as cardiac fibrogenesis (Działo, Tkacz, and Błyszczuk 2018), tumor-cell growth (Alkasalias et al. 2017; Datta et al. 2017b; Lin and Zheng 2015) or pulmonary artery hypertension (MacKay et al. 2017; Wei et al. 2019; Zhang and Wu 2017). The cellular effect observed in fibroblast activation is associated to changes in cellular function and behaviour. Actin polymerization regulates cell polarization, organization of adhesion structures and the generation of the force essential for cell migration (Smith and Barton 2018). These events allow cells to migrate into the injured site where they proliferate and release components of the ECM to regenerate the damaged tissue. However, an excessive activation of fibroblasts increases the ECM deposition leading to accumulated fibrogenesis (Ieronimakis et al. 2016). Although the actin rearrangement produced in fibroblast activation has not been characterized in FAPs, previous studies suggest that fibroblasts and FAPs are phenotypically and biochemically equivalent supporting that Rho-kinase activation could have a similar effect in both cells (Contreras, Rossi, and Brandan 2019). We demonstrated that RhoA/ROCK2 pathway is increased in muscles of DMD patients when compared to age and sex matched healthy controls subjects. Therefore, we decided to analyse the effect of PDGF-AA on RhoA/ROCK2 signalling pathway in FAPs isolated from DMD patients. First, we confirmed that RhoA/ROCK2 pathway can be modulated by PDGF-AA since C3-exoenzyme and fasudil significantly reduced the signalling pathway activation. Our results show that PDGF-AA activated RhoA/ROCK2 pathway resulting in an increase of actin filament polymerization, collagen release *in vitro* and proliferation and migration and collagen release *in vitro* and that both C3 and fasudil attenuated these effects. Although PDGF-AA induced a higher proliferation rate at 48 and 72 hours, the blocking effect of RhoA/ROCK2 inhibitors was only observed at 72 hours. The lack of proliferation inhibition at 48 hours after C3-exoenzyme or fasudil treatment could be due to other signalling pathways activated after PDGF-AA treatment. PDGF-AA is a mitogen growth factor that not only activates RhoA pathway but also triggers other signalling pathways, such as MAPK, PLC or PI3K. While these molecular pathways are mainly involved in cell proliferation, RhoA/ROCK2 is mainly involved in actin remodelling.

Since the Rho/ROCK-mediated pathway interact with other signaling pathways known to contribute to fibrosis (Bonniaud et al. 2005; Denton and Abraham 2001; Watts and Spiteri 2004), we tested whether fasudil reduces fibrosis in the dba/2J-*mdx* mice. Fasudil is a highly selective inhibitor of ROCK1 and ROCK2 isoforms that has been used in the clinic as a first-generation selective Rho/ROCK inhibitor (Xu et al. 2017). ROCK polymerization of actin in different cells leading to different effects such as control of cell size, migration, modulation of gene expression, differentiation or cell transformation. Fasudil has been tested as a therapeutic agent for different diseases as pulmonary hypertension, amyotrophic lateral sclerosis or cardiovascular disease (Koch et al. 2020; Xie et al. 2018). We observed that dba/2J-mdx mice treated with fasudil had better performance on grip strength test than non-treated mice. These results were associated to a decrease in the amount of collagen-I and in the number of FAP cells in the skeletal muscles of the treated animals compared to the non-treated ones. Our findings suggest that treatment with fasudil may reduce fibrosis through blocking FAPs proliferation. These results are in accordance with previous studies using fasudil as an antifibrotic agent. Baba et al. studied the mechanisms underlying renal interstitial fibrosis induced by unilateral uretral obstruction in mice and found that fasudil reduced collagen area and macrophage infiltration in the affected kidney (Itsuko Baba, 2015). The effects of fasudil were also studied on idiopathic pulmonary fibrosis. Fasudil treatment on bleomycin-induced pulmonary fibrosis mice reduced the total content of collagen, the number of infiltrating macrophages and the production of TGF-β1, CTGF, alpha-smooth muscle actin (α-SMA), and plasminogen activator inhibitor-1 (PAI-1) in the lungs of treated animals ((Jiang et al. 2012). Qi XJ, 2015)(Li et al. 2018),

Since the inflammatory response is a characteristic feature of DMD muscles, we also analysed the profile of the macrophages present in skeletal muscle after treatment with fasudil. We observed that this drug induced several changes including a reduction in the number of macrophages and a reduction in the number of CD163 and CD206 positive macrophages without modifying the M1 population. These results are in agreement with previous studies. For example, Xie et al. studied the effect of fasudil in preventing hepatic fibrosis in type 1 diabetic rats and identified a reduced inflammatory response in the liver of treated mice (Y. Xie, 2018). Our results can have two different explanations: 1) the attenuation in the fibrotic process in fasudil can improve muscle regeneration in treated animals preventing an increase of macrophages infiltrating the injured muscle or 2) as ROCK2 is also expressed in macrophages, the inhibition of this pathway reduced their migration into injured muscles. These findings are in accordance with other authors who studied ROCK2 as a modulator of M2 polarization. Zandi et al. demonstrated that ROCK isoforms present a dual effect on macrophage polarization. On the one hand, ROCK2 inhibition suppressed M2 and furthered M1 polarization, while ROCK1 inhibition furthered M2 polarization in age-related macular degeneration (Zandi et al. 2015). ROCK2 signalling has also been shown to induce M2 polarization in human primary monocytes(Lapointe et al. 2020). Although we did not observe an increase in M1 polarization, the inhibition of ROCK2 by fasudil presented a tendency towards a decreased expression of M2 macrophages markers. We think that the improved grip-strength test reduced expression of collagen-I together with the tendency towards a decreased number of PDGFRα+ cells on skeletal muscles in fasudil treated mice could be due to: 1) RhoA/ROCK2 inhibition has a direct effect on fibrosis deposition by decreasing the proliferation of FAPs in the muscle and 2) the decreased inflammatory response and decreased M2 polarization after fasudil treatment positively contributes to maintain an anti-inflamatory microenvironment that will involve a decreased FAPs activation together with a higher regeneration of the muscle.

Although ROCK is expressed in different cell types, the side effects produced by fasudil does not include major safety concerns although it can produce allergic skin reaction, reversible renal impairment or a slight drop in systolic blood pressure (Koch et al. 2020). Fasudil targets the ATP-dependent kinase domain of either ROCK1 and ROCK2 with equal potency and without selective effects. It is worth noting that ROCK2 is the isoform detected in our proteomic analysis and previous studies have shown that ROCK2 is the predominant isoform in skeletal muscle(Pelosi et al. 2007). Since we have not observed side effects on the dba-2J/mdx mice, the use of a selective ROCK2 inhibitor could be further studied as a higher selective drug for muscle fibrotic development (Boerma et al. 2008).

The fibrotic process that occurs in muscle dystrophies is an irreversible process that should be prevented before the accumulation of ECM components disturbs the muscle architecture and function. Since FAP activation and ECM remodelling start early during skeletal muscle degeneration, inhibiting the initial cellular changes that occur on FAPs could be targeted to prevent the increase of ECM deposition components. Therapies focused on targeting actin rearrangement to prevent FAP migration into the injury site could help to slow down the progression of the fibrogenic process due to these events will be followed by an increase in FAP proliferation and differentiation Coupling anti-fibrotic therapies with cell therapy or gene therapy could attain a better outcome of the therapy. Then, by inhibiting the ROCK2 signalling in FAPs we are targeting the first stages of fibrosis which could be further studied as a potential treatment to attenuate the disease progression and muscle fibrosis in DMD patients.

In summary, the present study demonstrates that PDGF-AA induces RhoA/ROCK2 pathway signalling in DMD FAPs leading to their proliferation, migration and production of components of the ECM, including collagen-I. Treatment with fasudil, a well-known ROCK inhibitor, blocks the effect of PDGF-AA on cells *in vitro* and reduces muscle fibrosis increasing muscle strength in a DMD murine model. Our results open the door to new studies further characterizing the efficacy of more specific ROCK2as potential new treatments for DMD.

## FIGURE LEGENDS

**Supplemental 1.**
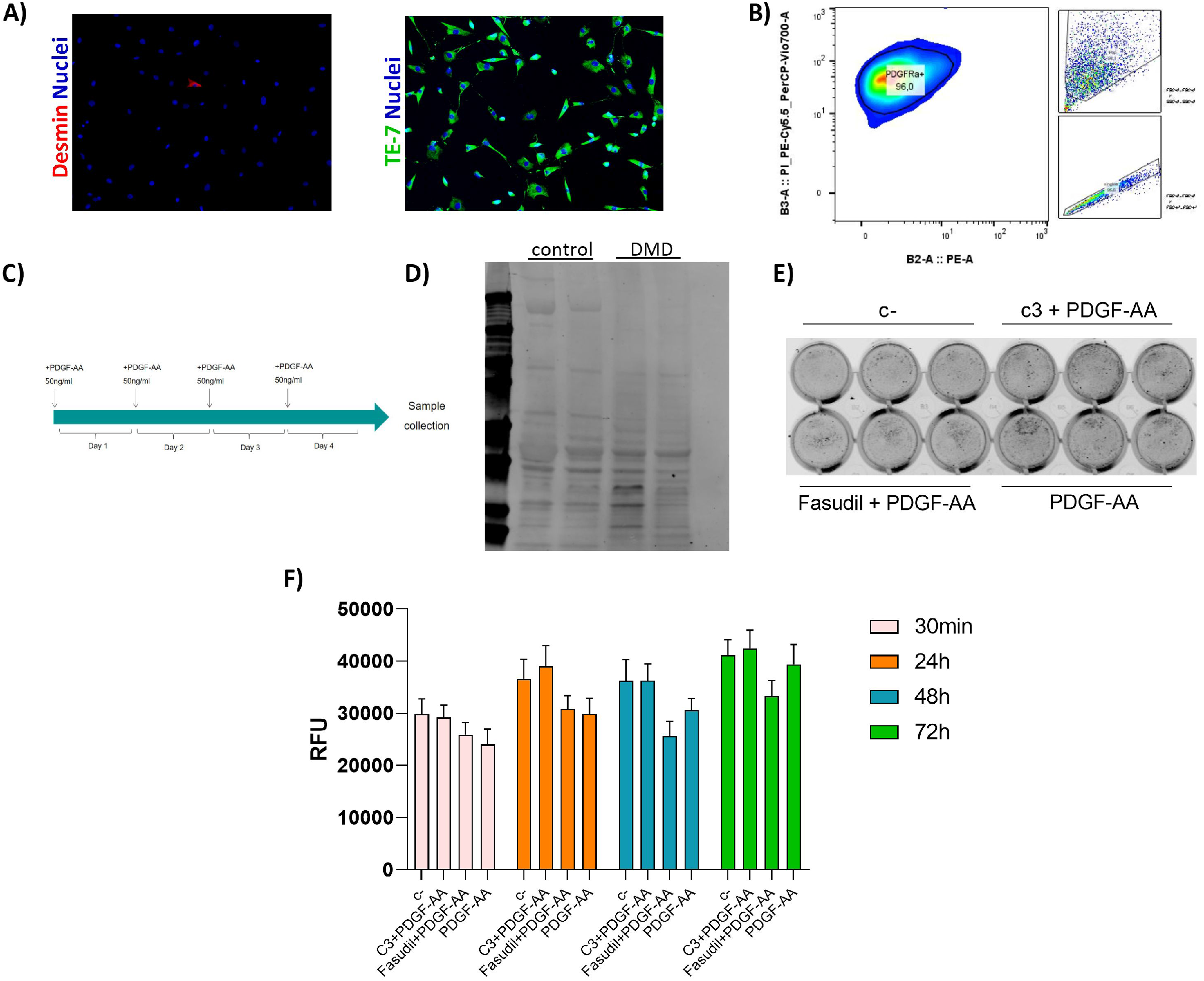
A) Representative images of the DMD cells. CD56-were stained with desmin and TE-7 markers. B) Analysis of the PDGFRα in the CD56-isolated cells showing positivity for the receptor. C) Experimental plan to screen. D) Representative WB of total protein bands measured with the Revert 700. E) Representative in-cell western image showing the total cell number signal stained with TO-PRO-3. F) Viability determination using the PrestoBlue cell viability assay to compare the effect of PDGF-AA, C3 and fasudil on FAPs at 30min, 24h, 48h and 72h.Data are represented as the mean of 3 replicates ± standard error of the mean. Results were statistically analysed using one-way ANOVA followed by Tukey post hoc test.

**Supplemental 2.**
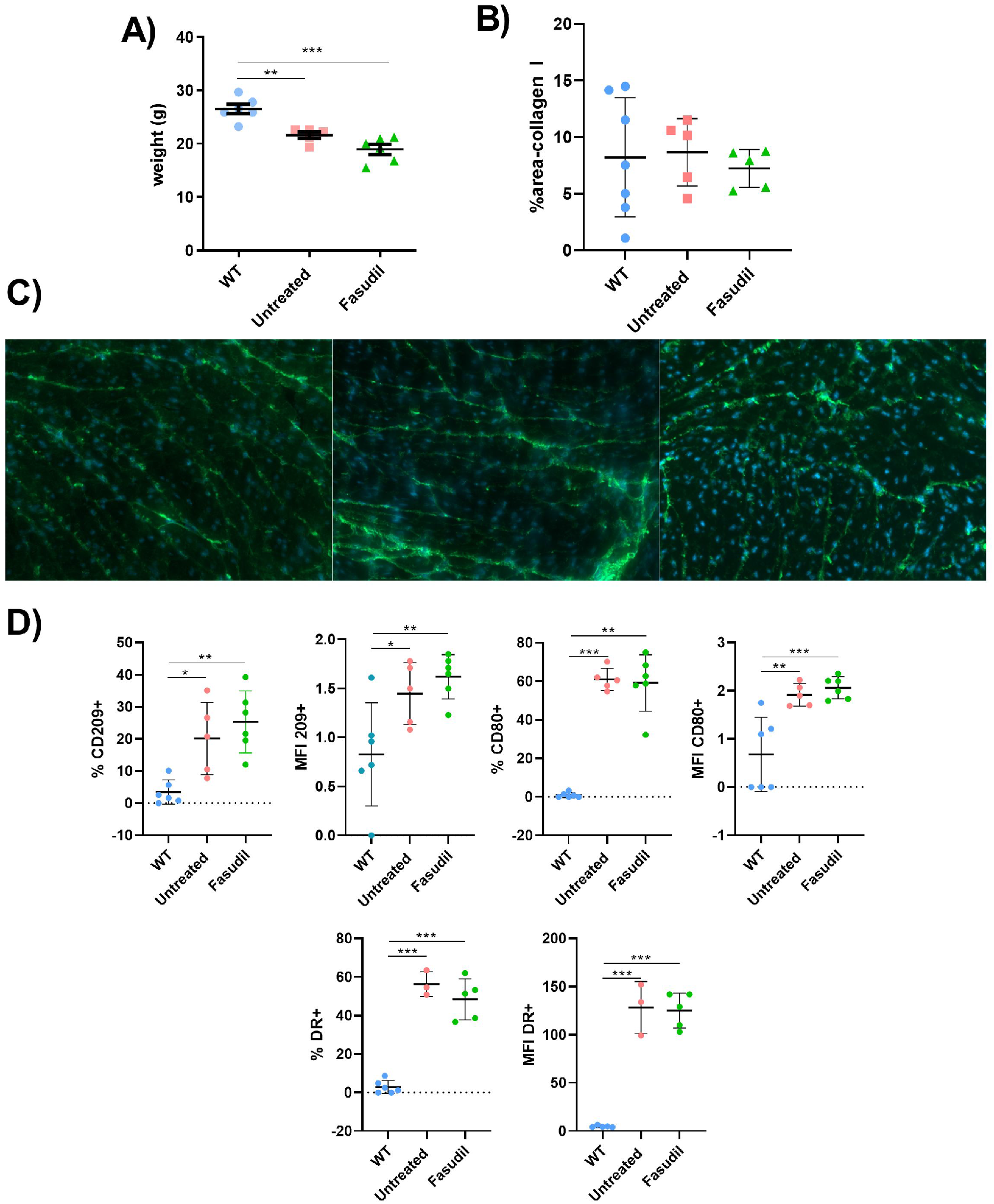
A) Effect of fasudil treatment in the dba/2J-*mdx* mice.A) Weight measurement after 6 weeks treatment with fasudil in dba-2J WT mice, untreated dba/2J-*mdx* and fasudil treated dba/2J-*mdx*. B) Quantification of collagen I expression area in the heart E) Representative immunofluorescence images of collagen I expression in the heart. Immunofluorescence images were taken at 10X magnification. D)Percentage and mean intense fluorescence (MFI) of CD209, CD80 and DR markers analysed by flow cytometry. Data are represented as the mean of the groups analysed ± standard error of the mean. Results were statistically analysed using one-way ANOVA followed by Tukey post hoc test. Statistical significance was set at p<0.05. *p<0.05; **p<0.01; ***p<0.0001.

## ACKNOWLEDGMENTS

This study has been funded by Instituto de Salud Carlos III through the project “PI18/01525” (Co-funded by European Union ERDF), by AFM (France) through the project 22525 and by Fundación Isabel Gemio. XSC was supported by a “Sara Borrell” post-doctoral fellowship (CD18/00195), Fondo Social Europeo (FSE), Instituto de Salud Carlos III (Spain).

## COMPETING INTERESTS

The authors declare that they have no competing interests.

## REFERENCES

Alkasalias, Twana et al. 2017. “RhoA Knockout Fibroblasts Lose Tumor-Inhibitory Capacity in Vitro and Promote Tumor Growth in Vivo.” Proceedings of the National Academy of Sciences of the United States of America 114(8): E1413–21.

Andrae, Johanna, Radiosa Gallini, and Christer Betsholtz. 2008. “Role of Platelet-Derived Growth Factors in Physiology and Medicine.” Genes and Development 22(10): 1276– 1312.

Birnkrant, J et al. 2018. “Diagnosis and Management of Duchenne Muscular Dystrophy, Part 2: Respiratory, Cardiac, Bone Health, and Orthopaedic Management.” The Lancet Neurology 17(18): 347–61.

Bonniaud, Philippe et al. 2005. “TGF-β and Smad3 Signaling Link Inflammation to Chronic Fibrogenesis.” The Journal of Immunology 175(8): 5390–95.

Bushby, Katharine et al. 2010. “Diagnosis and Management of Duchenne Muscular Dystrophy, Part 1: Diagnosis, and Pharmacological and Psychosocial Management.” The Lancet Neurology 9(1): 77–93.

Chen, Xuesong, Kristin Pavlish, and Joseph N. Benoit. 2008. “Myosin Phosphorylation Triggers Actin Polymerization in Vascular Smooth Muscle.” American Journal of Physiology - Heart and Circulatory Physiology 295(5): H2172.

Contreras, Osvaldo, Fabio M. Rossi, and Enrique Brandan. 2019. “Adherent Muscle Connective Tissue Fibroblasts Are Phenotypically and Biochemically Equivalent to Stromal Fibro/Adipogenic Progenitors.” Matrix Biology Plus 2:100006

Datta, Anirban, Emma Sandilands, Keith E. Mostov, and David M. Bryant. 2017a. “Fibroblast-Derived HGF Drives Acinar Lung Cancer Cell Polarization through Integrin-Dependent RhoA-ROCK1 Inhibition.” Cellular Signalling 40: 91–98.

Denton, Christopher P., and David J. Abraham. 2001. “Transforming Growth Factor-β and Connective Tissue Growth Factor: Key Cytokines in Scleroderma Pathogenesis.” Current Opinion in Rheumatology 13(6): 505–11.

Działo, Edyta, Karolina Tkacz, and Przemysław Błyszczuk. 2018. “Crosstalk between the TGF-β and WNT Signalling Pathways during Cardiac Fibrogenesis.” Acta Biochimica Polonica 65(3): 341–49.

Fabregat, Antonio et al. 2018. “Reactome Graph Database: Efficient Access to Complex Pathway Data.” PLoS Computational Biology 14(1).

Fiore, Daniela et al. 2016. “Pharmacological Blockage of Fibro/Adipogenic Progenitor Expansion and Suppression of Regenerative Fibrogenesis Is Associated with Impaired Skeletal Muscle Regeneration.” Stem Cell Research 17(1): 161–69.

Ge, Zhengxing et al. 2016. “Basic Fibroblast Growth Factor Activates b -Catenin / RhoA Signaling in Pulmonary Fibroblasts with Chronic Obstructive Pulmonary Disease in Rats.” Molecular and Cellular Biochemistry 423(1-2): 165–174.

Hogarth, Marshall W. et al. 2019. “Fibroadipogenic Progenitors Are Responsible for Muscle Loss in Limb Girdle Muscular Dystrophy 2B.” Nature Communications 10(1): 2430.

Ieronimakis, Nicholas et al. 2016. “PDGFRα Signalling Promotes Fibrogenic Responses in Collagen-Producing Cells in Duchenne Muscular Dystrophy.” The Journal of pathology 240(4): 410–24.

Jiang, Chunguo et al. 2012. “Fasudil, a Rho-Kinase Inhibitor, Attenuates Bleomycin-Induced Pulmonary Fibrosis in Mice.” International Journal of Molecular Sciences 13(7): 8293– 8307.

Joe, Aaron W B et al. 2010. “Muscle Injury Activates Resident Fibro/Adipogenic Progenitors That Facilitate Myogenesis.” Nature Cell Biology 12(2):153–63.

Kazlauskas, Andrius, and Jonathan A. Cooper. 1989. “Autophosphorylation of the PDGF Receptor in the Kinase Insert Region Regulates Interactions with Cell Proteins.” Cell 58(6): 1121–33.

Kelly, James D et al. 1991. “Platelet-Derived Growth Factor (PDGF) Stimulates PDGF Receptor Subunit Dimerization and Intersubunit Trans-Phosphorylation”. Journal of Biological Chemistry 266(14): 8987–92.

Kharraz, Yacine et al. 2014. “Understanding the Process of Fibrosis in Duchenne Muscular Dystrophy.” BioMed Research International 965631.

Klingler, Werner, Karin Jurkat-Rott, Frank Lehmann-Horn, and Robert Schleip. 2012. “The Role of Fibrosis in Duchenne Muscular Dystrophy.” Acta myologica : myopathies and cardiomyopathies : official journal of the Mediterranean Society of Myology 31(3): 184– 95.

Koch, Jan C. et al. 2020. “Compassionate Use of the ROCK Inhibitor Fasudil in Three Patients With Amyotrophic Lateral Sclerosis.” Frontiers in Neurology 11: 173.

Lapointe, Fanny et al. 2020. “RPTPε Promotes M2-Polarized Macrophage Migration through ROCK2 Signaling and Podosome Formation.” Journal of Cell Science 133(5): 234641.

Li, Xiao-Dong et al. 2018. “Fasudil Inhibits Actin Polymerization and Collagen Synthesis and Induces Apoptosis in Human Urethral Scar Fibroblasts via the Rho/ROCK Pathway.” Drug design, development and therapy 12: 2707–13.

Liebermeister, Wolfram et al. 2014. “Visual Account of Protein Investment in Cellular Functions.” Proceedings of the National Academy of Sciences of the United States of America 111(23): 8488–93.

Lin, Yuan, and Yi Zheng. 2015. “Approaches of Targeting Rho GTPases in Cancer Drug Discovery.” Expert opinion on drug discovery 10(9): 991–1010.

MacKay, Charles E et al. 2017. “ROS-Dependent Activation of RhoA/Rho-Kinase in Pulmonary Artery: Role of Src-Family Kinases and ARHGEF1.” Free radical biology & medicine 110: 316–31.

Makino, Katsunari et al. “Blockade of PDGF Receptors by Crenolanib Has Therapeutic Effect in Patient Fibroblasts and in Preclinical Models of Systemic Sclerosis.” The Journal of Investigative Dermatology 137(8): 1671–1681.

Mann, Christopher J. et al. 2011. “Aberrant Repair and Fibrosis Development in Skeletal Muscle.” Skeletal Muscle 1(1):21.

Olson, Lorin E, and Philippe Soriano. 2009. “Increased PDGFRalpha Activation Disrupts Connective Tissue Development and Drives Systemic Fibrosis.” Developmental cell 16(2): 303–13.

Pelosi, Michele et al. 2007. “ROCK2 and Its Alternatively Spliced Isoform ROCK2m Positively Control the Maturation of the Myogenic Program.” MOLECULAR AND CELLULAR BIOLOGY 27(17): 6163–76.

Perandini, Luiz Augusto, Patricia Chimin, Diego da Silva Lutkemeyer, and Niels Olsen Saraiva Câmara. 2018. “Chronic Inflammation in Skeletal Muscle Impairs Satellite Cells Function during Regeneration: Can Physical Exercise Restore the Satellite Cell Niche?” The FEBS Journal 285(11): 1973–84.

Phelps, Michael, Pascal Stuelsatz, and Zipora Yablonka-Reuveni. 2016. “Expression Profile and Overexpression Outcome Indicate a Role for ?Klotho in Skeletal Muscle ibro/Adipogenesis.” The FEBS journal 283(9): 1653–68.

Piñol-Jurado, Patricia et al. 2018. “Nintedanib Decreases Muscle Fibrosis and Improves Muscle Function in a Murine Model of Dystrophinopathy.” Cell death & disease 9(7): 776.

Schindelin, Johannes et al. 2012. “Fiji: An Open-Source Platform for Biological-Image Analysis.” Nature methods 9(7): 676–82.

Scotton, Chris J., and Rachel C. Chambers. 2007. “Molecular Targets in Pulmonary Fibrosis: The Myofibroblast in Focus.” Chest 132(4): 1311–21.

Shang, Xun et al. “Small-Molecule Inhibitors Targeting G-Protein-Coupled Rho Guanine Nucleotide Exchange Factors.” Proceedings of the National Academy of Sciences of the United States of America 110(8): 3155–60.

Smith, Lucas R., and Elisabeth R. Barton. 2018. “Regulation of Fibrosis in Muscular Dystrophy.” Matrix Biology 68–69: 602–15.

Svegliati, Silvia et al. 2007. “Stimulatory Autoantibodies to PDGF Receptor in Patients with Extensive Chronic Graft-versus-Host Disease.” Blood 110(1): 237–41.

Uezumi, Akiyoshi et al. 2011. “Fibrosis and Adipogenesis Originate from a Common Mesenchymal Progenitor in Skeletal Muscle.” Journal of cell science 124(21): 3654–64.

Wallace, Gregory Q., and Elizabeth M. McNally. 2009. “Mechanisms of Muscle Degeneration, Regeneration, and Repair in the Muscular Dystrophies.” Annual Review of Physiology 71(1): 37–57. 4, 2020).

Watts, Keira L., and Monica A. Spiteri. 2004. “Connective Tissue Growth Factor Expression and Induction by Transforming Growth Factor-β Is Abrogated by Simvastatin via a Rho Signaling Mechanism.” American Journal of Physiology-Lung Cellular and Molecular Physiology 287(6): L1323–32.

Wei, Hongwei et al. 2019. “Rho Signaling Pathway Enhances Proliferation of PASMCs by Suppressing Nuclear Translocation of Smad1 in PAH.” Experimental and therapeutic medicine 17(1): 71–78.

Xie, Y et al. 2018. “Fasudil Alleviates Hepatic Fibrosis in Type 1 Diabetic Rats: involvment of the inlammation and RhoA/ROCK pathway.” European Review for Medical and Pharmacolofical Sciences 22(17): 5665–5677.

Xu, Ning et al. 2017. “Fasudil Inhibits Proliferation and Collagen Synthesis and Induces Apoptosis of Human Fibroblasts Derived from Urethral Scar via the Rho/ROCK Signaling Pathway.” American journal of translational research 9(3): 1317–25.

Zandi, Souska et al. 2015. “ROCK-Isoform-Specific Polarization of Macrophages Associated with Age-Related Macular Degeneration.” Cell Reports 10(7): 1173–86.

Zhang, Yiqing, and Shangjie Wu. 2017. “Effects of Fasudil on Pulmonary Hypertension in Clinical Practice.” Pulmonary Pharmacology and Therapeutics 46: 54–63.

Zhao, Yajuan et al. 2003. “Platelet-Derived Growth Factor and Its Receptors Are Related to the Progression of Human Muscular Dystrophy: An Immunohistochemical Study.” The Journal of Pathology 201(1): 149–59.

